# Applying the area fraction fractionator (AFF) probe for total volume estimations of somatic, dendritic and axonal domains of the nigrostriatal dopaminergic system in a murine model

**DOI:** 10.1101/2023.10.19.562678

**Authors:** Alejandro Oñate-Ponce, Catalina Muñoz, Alejandra Catenaccio, Felipe A. Court, Pablo Henny

## Abstract

Using the nigrostriatal dopaminergic system as a case example, we describe a protocol based on Cavalieri’s principle to estimate the total volume of its somatic, nuclear, dendritic and axonal domains in the adult mouse. This protocol requires serial section analysis, a confocal laser scanning microscope (CLSM), and a stereological software combining the serial section manager (SSM) tool and the area fraction fractionator (AFF) probe. The protocol consists of the following steps: 1) systematic random sampling (SRS) of sites across sections for relevant regions of interest (ROI), which in this study included the substantia nigra pars compacta (SNc) and pars reticulata (SNr), the nigrostriatal tract, and the caudate-putamen complex (CP), 2) high resolution image acquisition of all SRS sites within and across sections, 3) loading of images on the previously defined SRS sites, 4) marking of all cellular structures of interest (cSOI) in the images, including perikarya and cell nuclei, dendrites, axonal segments and varicosities, using Cavalieri’s point-counting grids, and 5) estimation of cSOI total volume using the estimated area of each cSOI in every brain section, and the distance between sections. The added volume for all cSOI comprising the nigrostriatal dopaminergic system, in this murine model is around 0.37 mm^3^, with approximately 5% corresponding to perikarya and cell nuclei, 10% to neuropil/dendrites in substantia nigra, and 85% to axonal segments and varicosities within the CP. The use of a simple quantitative approach to assess the global state of a system may allow quantification of compartment-specific changes that may accompany neurodegenerative processes.

## Introduction

Design-based stereological techniques allow quantitative descriptions of three-dimensional structures and population size. These techniques make use of random and uniform sampling strategies, in conjunction with accurate geometric probes to eradicate systematic inaccuracies [1–5]. In neuroanatomical studies, stereological tools are applied to determine size of brain regions, as well as size and number of various brain cell types and other small nervous elements, under normal and experimental conditions [6–11].

### Cavalieri method and Area Fraction Fractionator

A stereological probe of widespread use is the Cavalieri method or estimator, which allows robust volume estimations [2–5, 9, 12]. As its name indicates, it is based on the Cavalieri principle that the volume of an object can be estimated from: 1) the cross sectional areas of an object as it is cut in equivalent and parallel sections and 2) the distance between sections [2]. The Cavalieri method was developed to include point-counting probes (i.e. a grid of points to mark whether the structure to quantify is present or not beneath points) to facilitate counting [1, 4, 5] and is routinely used for volume measurement in various brain regions, including hypothalamus, hippocampus, specific cortical areas, and whole brains [10, 11, 13, 14].

However, although widely used for volume estimation of brain regions, the Cavalieri method has not been applied to measure volume of small nervous elements, apart from two studies reporting its use for cell somata [15] or mitochondria [2] size. Indeed, cell volume is generally estimated using other stereological methods, such as the optical rotator or nucleator [1, 10, 16, 17].

A stereological probe derived from Cavalieri method is the Area Fraction Fractionator (AFF). The probe is carried out using a fractionator tool over the area of interest, so as to apply a point-counting grid at each sampling site, and not the entire region under study. This method has allowed efficient and robust area and number estimations of two dimensional elements, such as axons in nerve or tracts [3, 9, 18, 19]. It is, on the other hand, intriguing that the AFF probe has been mostly utilized for area estimations [9] and not, for instance, combined with serial sectioning analysis for estimations of small objects’ volume, as with the Cavalieri method [2].

### The nigrostriatal dopaminergic system: a case example

The nigrostriatal dopaminergic system is comprised by a population of dopamine-synthesizing and -releasing neurons, whose cell bodies mostly locate in the substantia nigra pars compacta (SNc) and project, through the nigrostriatal tract, to the caudate-putamen complex (CP) [20]. Here, axons bifurcate profusely [21] and release dopamine which in turn modulates activity in postsynaptic striatal neurons [20, 22]. The system plays an important role in motor control, most conspicuously evident in Parkinson’s disease, which is characterized by a progressive loss of nigrostriatal dopaminergic neurons that results in rigidity, bradykinesia and akinesia [23].

In this context, the relevance of accurate physical estimations of the several cellular and subcellular components of this system lies in the fact that changes in axonal, somatic, and dendritic structure can precede or accompany dysfunction of the nigrostriatal system occurring in PD or PD models [23–27].

In contrast to the vast amount of data on number of nigral dopaminergic cells [28, 29], dopaminergic synapses at the level of the striatum, or the volume of the substantia nigra or striatum [12, 30] there is less quantitative information about other physical and architectural dimensions of the intact dopaminergic system itself. Hence, the goal of this study was to generate a protocol that combines the aforesaid stereological Cavalieri principle and the AFF probe to provide an estimation of the total and relative volume of various elements of a neuronal system and to illustrate its application in the neuroanatomical description of the nigrostriatal dopaminergic system in the adult mouse.

To do that, we used the AFF probe in combination with serial section analysis. Because volume estimations depend on precise cross sectional area estimations, special emphasis was put on the use of high resolution and thin optical sections images through the use of the CLSM. We assessed the volume occupied by perikarya, cellular nuclei and neuropil (which is mostly comprised by dendrites, see Discussion) in the substantia nigra, axons in the nigrostriatal tract, and axonal varicosities and non-varicose axonal segments in the CP.

## Materials

### Animals and sample preparation

Brains of male adult mice were donated by Peter J. Magill and Paul D. Dodson (Oxford Parkinson’s Disease Centre). Mice were produced as part of Parkinson Disease model using a bacterial artificial chromosome (BAC) method in order to express human α-synuclein gene (SNCA) throughout dopamine neurons in the mice [31]. In this case, we worked with the hα-syn transgenic line which was originally used as the control group due to its low to mild levels of the expression of human α-syn, no dopaminergic cell loss during aging and, as compared to WT mice, normal number of SNc dopaminergic neurons and behavior [31]. All experimental procedures involving these live mice were conducted in accordance with the Animals (Scientific Procedures) Act, 1986 (United Kingdom).

Mice were perfused trans-cardially utilizing PBS (pH 7.4) and 4% paraformaldehyde. The brains were post-fixed in 4% paraformaldehyde for 12 hours at room temperature. Then, they were cryoprotected in 30% sucrose until they submerged in the solution, frozen and sent overseas. They were then cut in 25 μm thick coronal sections, employing a freezing-stage microtome. The brain sections were systematically collected in 8 series and hoarded in PBS at 4° C.

### Immunofluorescence

To characterize the dopaminergic system we utilized immunostaining for tyrosine hydroxylase (TH) [32, 33]. Coronal sectioned adult brains were used with the aim of observing the expression of this enzyme on diverse neuronal structures, such as perikaryon, neuropil, and axons. TH staining also allowed the identification of dopaminergic cells nuclei, which do not stain for TH but are surrounded by a TH positive perikaryon. For immunofluorescence, the sections were rinsed in PBS and blocked in 3% normal horse serum solution in PBS (Jackson ImmunoResearch) for 2 hours at room temperature. Subsequently, the sections were washed in PBS for 10 min and incubated with guinea pig anti-TH antibody (1/1000, Synaptic System) in a blocking solution for 12 hours. The sections were then rinsed in PBS and incubated for 2 hours in Alexa Fluor 488-conjugated donkey anti-guinea pig antibody (1/100, Jackson ImmunoResearch) diluted in blocking solution and washed in PBS. All the sections were mounted on glass slides in Vectashield mounting medium (Vectashield Vector Laboratories, Burlingame, CA).

A series of three additional adult mice animals were used, in this case for the specific analysis of the nigrostriatal tract (see below). The research protocol no. 08–2016 was approved by the Animal Care and Use Scientific Ethic Committee of Universidad Mayor. These transgenic mice were also generated as part of a Parkinson Disease model which evaluates the influence of RIPK3 ablation on the survival of dopaminergic neurons following a neurotoxic lesion in the striatum [34]. In this case, we analyzed the nigrostriatal tract on the control, non-lesioned hemisphere. Processing of brain and brain sections for TH immunohistochemistry was similar to the protocol described above except for primary (rabbit anti-TH antibody 1/200, Millipore ab152) and secondary (Alexa Fluor 555-conjugated donkey anti-rabbit, 1/500, Invitrogen A32794) antibody choice, and is reported in detail in [34].

### Microscopy, imaging and stereological probes

To image brain sections at low magnification an epifluorescence microscope (Nikon Eclipse Ci, Tokio Japan) equipped with a motorized x-y-z stage was used. For high magnification images, a CLSM (Zeiss LSM 700, Thornwood, NY) equipped with a 63X oil/water immersion objective (1.4 NA) for SNc, SNr and CP analysis and a CLSM (Sp8 Leica) with a 63x oil immersion objective (1.4 NA) for nigrostriatal tract analysis were used. Filters, pinhole size and emission / detection settings were to optimize vertical resolution, as based on Alexa Fluor 488 or 555 signals. Stereological probes were run on a PC utilizing the Stereo Investigator software (MBF Bioscience, 2019 and previous versions).

## Methods

The brain is cut in serial sections. The researcher needs to select the section cut thickness (this must be a uniform one), and the periodicity in which sections will be collected in one series. A further requirement to ensure a systematic random sampling approach is that the series to be worked with must be selected randomly (Figure 1).

**Figure 1.**
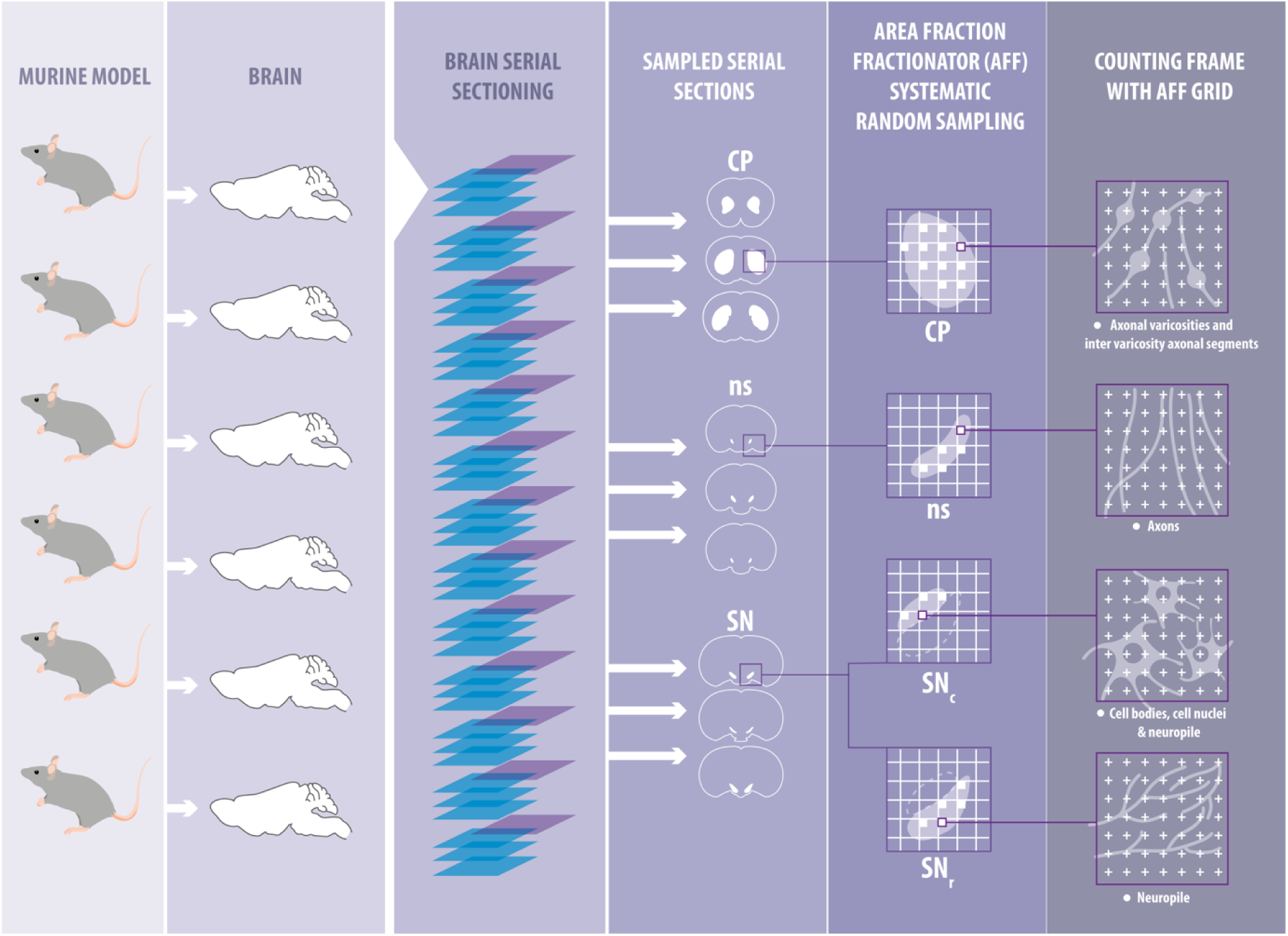
Protocol synopsis. Brains obtained from animals models under study (columns A and B) are collected in series (four in this example, column C). A randomly chosen series to work with is selected (in this case series 1, magenta, column C) and processed for identification of cellular structures of interest (cSOI, see column F). Regions of interest (ROIs, in this example SNc, SNr, nigrostriatal tract and CP) are analyzed across all the sections present throughout the anteroposterior axis (column D). Then, using the AFF probe, systematic random sampling sites are selected (column E) for imaging at high magnification. Images are taken for every sampling site, with a XY size that precisely matches the size of sampling sites (column F). These images are then analyzed using point-counting for area estimation of cSOI (column F).

We chose to work with the nigrostriatal dopaminergic system and utilized immunofluorescence staining for TH. TH staining allows a clear definition of the relevant regions of interest (ROI) in this system (see Figure 2). At the optical level, TH staining provides a signal that fills homogeneously the cytoplasmic compartment of the neuron’s different neuronal domains, and thus facilitates the identification and point-counting of positive structures (Figures 3 and 4).

**Figure 2.**
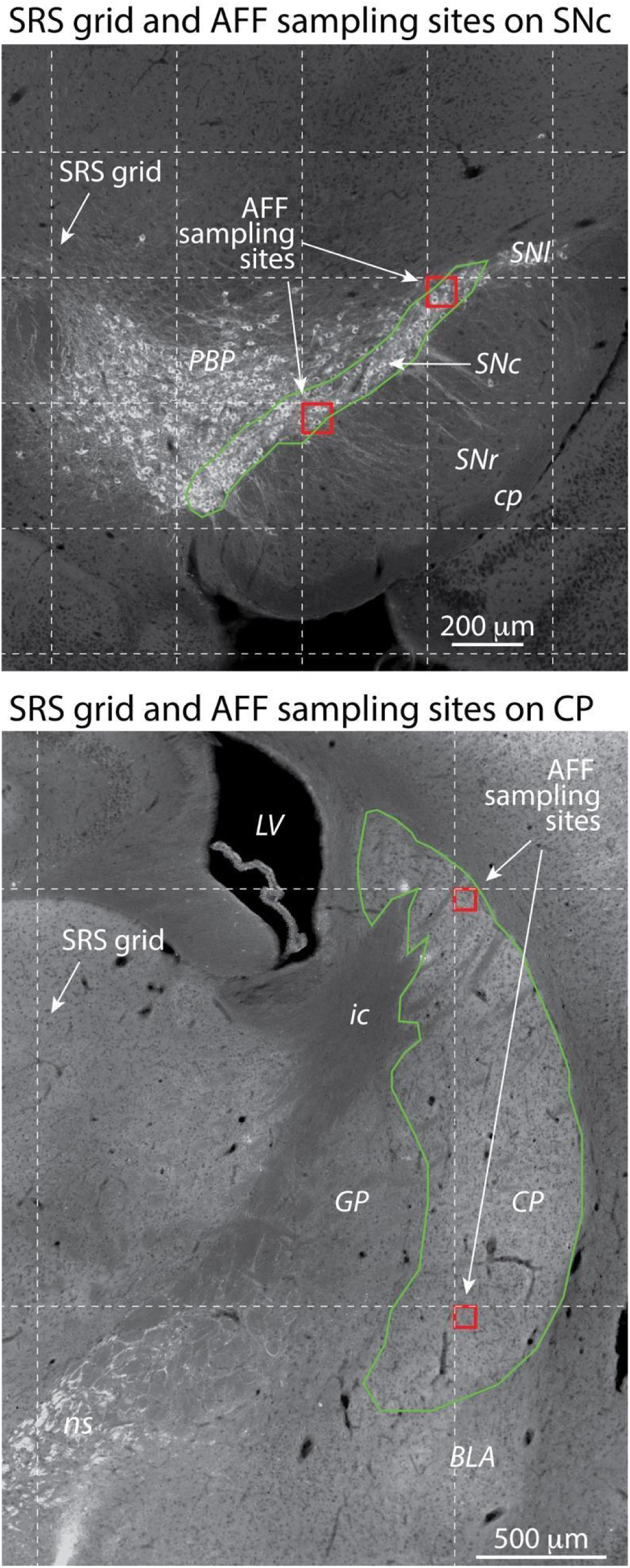
AFF systematic random sampling (SRS) grid and AFF sampling sites on CP and SNc. Top: Low magnification image of the SNc, with SRS grid (white grid, dashed lines), sampling sites (small red squares) and ROI contour (white line defining the SNc limits) overlaid. Grid for the SNc region was 350 μm x 350 μm. Sampling size on each sampling site was 79 μm x 79 μm. Because the SRS grid overlay position is defined randomly, so are the sampling sites and the portion of ROI that is sampled. Sampling sites are then acquired at high resolution. Bottom: Low magnification image of the CP, with SRS grid (white grid, dashed lines), sampling sites (small red squares) and ROI contour (white line defining the CP limits) overlaid. The SRS grid for the CP region was 1600 μm x 1600 μm. Sampling size on each sampling site was 79 μm x 79 μm. Anatomical abbreviations: BLA, basolateral amygdala; cp, cerebral peduncle; CP, caudate-putamen; GP, globus pallidus; ic, internal capsule; LV, lateral ventricle; ns, nigrostriatal tract; PBP, parabrachial pigmented nucleus of the VTA; SNc, SNl, SNr: substantia nigra pars compacta (c), lateralis (l) and reticulata (r).

**Figure 3.**
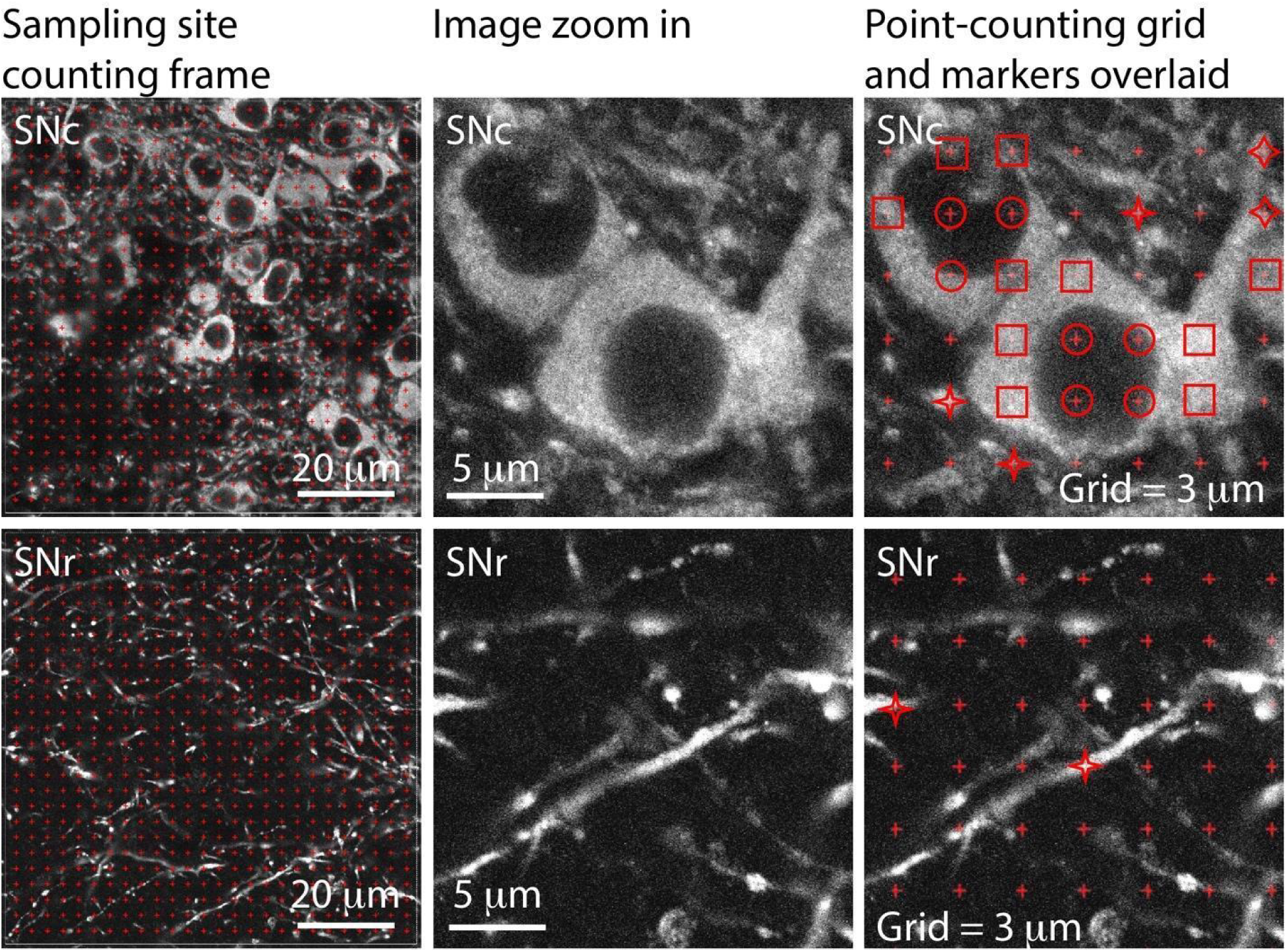
AFF sampling site on SNc and SNr, labeled for TH. AFF sampling sites are taken at high resolution with a CLSM and then opened in the AFF probe. There, point-counting is carried out for marking of cellular structures of interest (cSOI). Left column: an overall view of the AFF sampling site counting frame and point-counting grid (red crosses). Middle and right columns: zoom in showing the cSOI without (middle) and with (right) the point-counting grid. Here, markers for neuropil (four-point stars), perikaryon cytoplasm (squares) and cell nuclei (circles) are placed on the point-counting grid. Grid of points is 3 μm for both structures.

**Figure 4.**
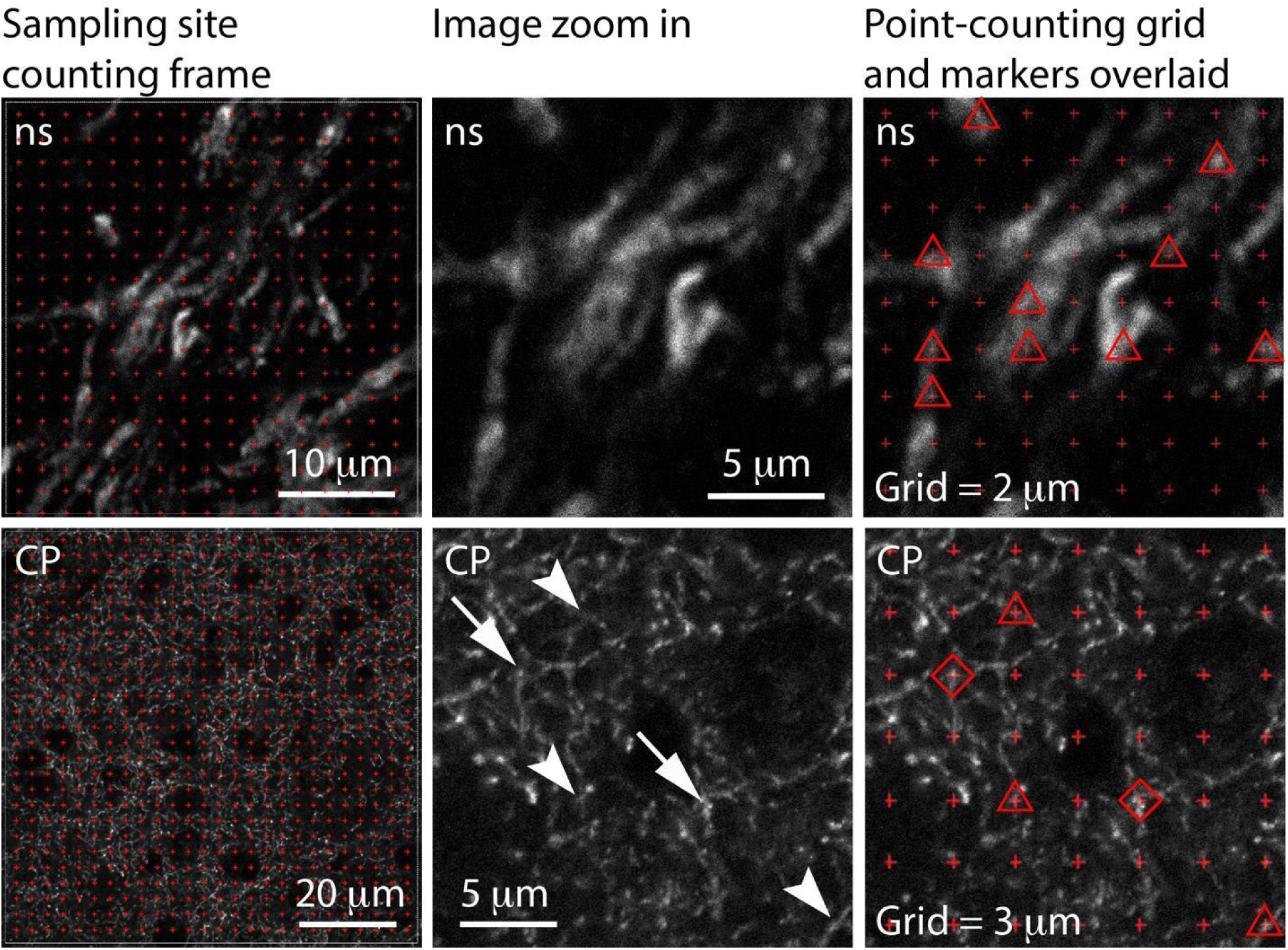
AFF sampling site on nigrostriatal tract and CP, labeled for TH. AFF sampling sites are taken at high resolution with a CLSM and then opened in the AFF probe. There, point-counting is carried out for marking of cellular structures of interest (cSOI). Left images: an overall view of the AFF sampling site counting frame and point-counting grid (red crosses). Middle and right columns: zoom in images with (middle) and without (right) the point-counting grid. Spacing between points is 2 μm for nigrostriatal tract (ns) structures (top right image) and 3 μm for CP structures (bottom right image). In the middle bottom image, arrows indicate axonal varicosities and arrowheads indicate axonal segments. In the right image, markers for axonal segments (triangles in nigrostriatal tract and CP) and axonal varicosities (rhomboids on CP) are placed on the point-counting grid. In the case of the right bottom, markers also correspond to arrows and arrowheads of the middle image.

The ROI contours must be defined in every section based on TH staining plus aid of a brain atlas and other criteria for definition of their limits. In our case, SNc and SNr contours were defined according to TH staining and following limits provided in Franklin and Paxinos [35]. For the nigrostriatal tract, the contours were defined combining two criteria: 1) approximate location and shape of the nigrostriatal tract contour of Franklin and Paxinos [35], and 2) a region containing a bundle of compactly arranged TH axons and that usually provides a high intensity signal. We chose two criteria because the definition of the nigrostriatal tract limits is not obvious due to the existence of multiple TH labeled axons that run in parallel to the nigrostriatal tract, such as those of the medial forebrain bundle. As defined in the Franklin and Paxinos atlas, we only defined nigrostriatal tract contours between Bregma levels AP -1.06 to -2.80 mm. For the CP, contours were defined according to Franklin and Paxinos, except for its ventral limit which in most levels is continuous with the nucleus accumbens. For that, we defined the ventral limit of the CP as an imaginary straight line running from dorsolateral end of the piriform cortex’s pyramidal layer towards the ventral end of the lateral ventricle between Bregma AP 1.78 to 0.74 mm. From there and towards more posterior levels, the contour was based on the atlas.

ROI contours are traced at low magnification and placed on the virtual serial section manager (SSM) in the Stereo Investigator software. This can be done online if the Stereo Investigator station is connected to a fluorescent microscope, or offline if low magnification images are taken in another microscope and imported. Then, a systematic random sampling (SRS) strategy to define sampling sites within each contour ROI is performed, in this case within the AFF probe. We defined the size of the SRS grid (i.e. the distance between the sampling sites, Figure 1) to be different for each ROI, with larger ROIs having larger SRS grid sizes, so as to reduce oversampling in larger ROI. Thus, chosen SRS grid sizes were 350 μm x 350 μm for SNc, 225 μm x 225 μm for nigrostriatal tract, 662 μm x 662 μm for SNr, and 1600 μm x 1600 μm for CP. Further AFF probe parameters, including the counting frame size (i.e. sampling site size) and the spacing between counting points in the counting frame are defined. A counting frame of 79 μm x 79 μm was defined for SNc, SNr and CP and a counting frame of 35 μm x 35 μm for nigrostriatal tract. Spacing between points in the counting frame grid was 3 μm for SNc, SNr and CP, 2 μm for the nigrostriatal tract. Parameter values are decided after several pilot probe runs in which SRS grid size, counting frame size, section interval, and point-counting grid spacing are tested and evaluated against Gundersen’s CE (m = 1) values. In most cases, sampling parameter values were considered adequate if obtained CE values were consistently below 0.1.

After the AFF probe has been started, a preview of the resulting SRS layout (including the SRS grid and the sampling sites) with the underlying low magnification ROI image (Figure 2) is screenshotted. AFF probes are started in all sections containing the ROI and therefore multiple screenshots are taken and printed. This facilitates the process of identifying sampling sites in the CLSM.

In the CLSM, counting frame images are taken at the same locations indicated by the SRS layout previously described. The size of the acquired image is the same as the counting frame size in order to optimize CLSM time use.

An important consideration on the practical use of the Cavalieri principle is the fact that images always have some degree of overprojection, which relates directly with the optical section thickness [3, 5]. It is critical then to ensure that images taken at sampling sites at high magnification are acquired with a high numerical aperture objective and wavelength-optimized pinhole size, so as to obtain the minimal optical section thickness [36].

Once images at every counting site at every section containing a ROI are taken, the previously saved AFF probe is opened again and images are loaded on the corresponding sampling sites (counting frames). Then, markers are used to indicate the presence of cSOI under the point-counting grid (Figure 3 and 4). We chose markers according to the cSOI predicted to be found in the different ROI of the nigrostriatal system. Thus, in the SNc we defined three types of cSOI: (1) perikarya, which more specifically correspond to the cell body cytoplasmic compartment, (2) cell nuclei, which correspond to the unlabeled compartment inside labeled perikarya, and (3) neuropil, which we defined as any structure different from a perikaryon or a cell nuclei, and that can be assumed to mostly comprise dendrites and a minor contingent of axons. We did not attempt to morphologically differentiate axons from dendrites, given the presence of varicose dendrites that may be confused with axons, as well as smooth and thin dendrites that may be otherwise taken for axons. In any case, given that axons from SNc neurons do not bifurcate as they advance towards the nigrostriatal tract (Henny et al., 2012), we assume that most of neuropil in the substantia nigra correspond to dendrites (Figure 3). In the SNr, we defined the same three types of cSOI, although they mostly corresponded to neuropil (Figure 3). In the nigrostriatal tract, we considered axonal segments as the only cSOI present. In the CP we identified axonal varicosities, which we defined as swelling and/or high intensity spots in the CP, and inter-varicose axonal segment, as those labeled structures that did not swell and usually connect two axonal varicosities (Figure 4).

For the sake of simplicity, we did not study other regions that could otherwise be considered part of the nigrostriatal dopaminergic system, including dopaminergic axons that branch off and innervate the subthalamic nucleus [37], or the axons running in the internal capsule as they reach the CP.

Following is a step by step summary of the procedure. This section should be taken as a framework that must be adapted to the circumstances and needs of the specific project carried out. As explained above, parameter values were obtained following multiple pilot testing.

### Step by step procedure

#### I. Systematic random sampling

1. Randomly chose a series to work with. In the present study, the brain was cut in 8 series and the specific series to be analyzed in each brain (1 to 8) was chosen randomly in www.random.org
2. Place the slide under the microscope at low magnification and image it in the Stereo Investigator software. Alternatively, take low magnification images on a different microscope (e.g. CLSM) and import them in Stereo Investigator.
3. Identify the first and last section of the series containing the region of interest.
4. Using the serial section manager (SSM) tool, define all the sections and respective top of section z-level containing the ROI. The interval value between the sections must be indicated. Although for AFF probes mounted thickness is not a feature that should directly or theoretically affect area or volume estimations, some value must be provided (e.g. the original cut thickness, 25 μm).
5. Delineate all ROI present in each section at low magnification using the SSM tool.
6. Go back to the first SSM section.
7. Define the size of the counting frame (CF). Note that this step must be carried out before starting the AFF.
8. Open the AFF probe and click on the ROI’s contour to be analyzed.
9. Define the SRS grid size, which corresponds to the distance between sampling sites (i.e. the counting frames).
10. Define the value of the counting frame point-counting grid spacing and accept changes.
11. Go to Preview SRS layout and then select Image/Fit to screen function so as to simultaneously see the low magnification image of the region to be analyzed, the ROI contour, the SRS grid and the SRS layout, with the sampling sites indicated. See Figure 2.
12. Print or take a screenshot.
13. Save the unfinished probe as a file that you can open later.

#### II. CLSM acquisition of sampling sites images

14. Place the slide under a low magnification objective and identify the section and ROI.
15. With the help of low magnification images and the SRS layout, find each of the sampling sites (SRS) on the ROI.
16. Move to a high magnification lens and acquire an image the size of the counting frame. Do this with the best possible optics, including best signal to noise ratio (which is usually achievable in the upper portion of the section where immunostaining may be best), appropriate pinhole for the emission wavelength and the emission/detection filter pair being used, minimal z-axis optical thickness, and maximal dynamic range for intensity.
17. Repeat procedure for all required sampling sites across sections.

#### III. Quantification

18. Back in the Stereo Investigator software, define a new lens to match the CLSM image resolution.
19. Open the previously saved file with the AFF probe.
20. Open the probe and go to the first sampling site.
21. Open the appropriate high magnification image taken at the CLSM and align it with the counting frame on screen.
22. Choose the markers for each of the cSOI you would like to estimate.
23. Mark the cSOI.
24. Move to the next sampling site (counting frame) and iterate until all sites in all sections have been sampled.
25. Once finished, check the results using Probeslll Probe run list.
26. Select all relevant section probes and view results.
27. Extract relevant data including number of markers, volume estimation per cSOI and sampling statistics such as Gundersen’s coefficient of error (CE).

### Case results

As shown in Table 1, the results of this example study indicate that the estimated volume of the entire nigrostriatal system of adult mice per hemisphere approximate 0.370 mm^3^. Up to 85% of this volume is located in the CP and corresponds both to axonal varicosities (48%) and inter varicosity axonal segments (37%). The remaining 15% corresponds to the cytoplasmic compartment of perikarya (3%), cell nuclei (2%) and neurites (10%) that are located in the substantia nigra. An almost negligible volume (less than 0.001 mm^3^) corresponds to the nigrostriatal tract. From our CE estimations, sampling was robust and within the values normally accepted for stereological estimates [7].

**Table 1.**
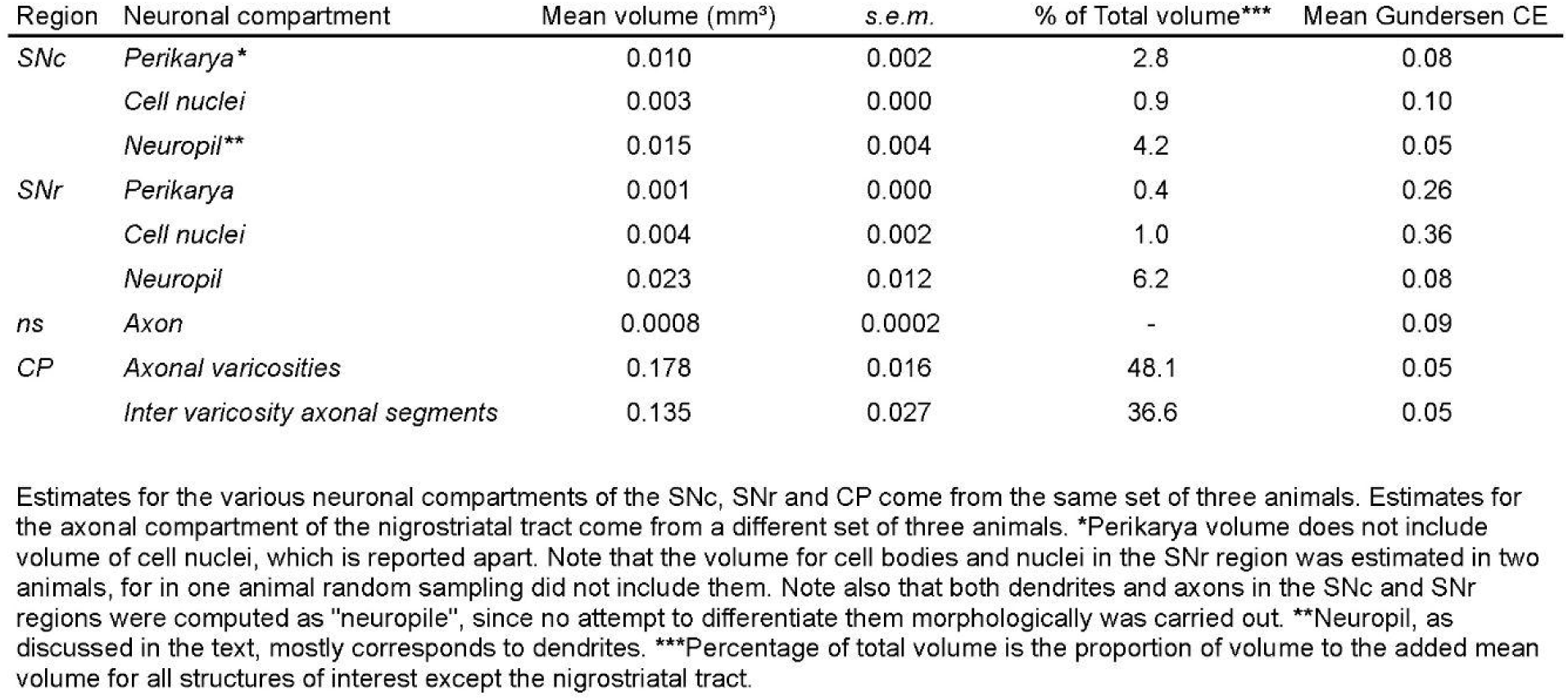
Volume estimates of neuronal compartments of the adult nigrostriatal dopaminergic system per hemisphere.

## Discussion

In this chapter, we described a protocol to estimate the entire volume of a specific neuronal system. We applied this protocol to the dopaminergic nigrostriatal system and reported that, in this specific model, the volume is about 0.37 mm^3^. The volume of the entire adult mouse brain has been estimated in 508 mm^3^ [38], thus indicating that the nigrostriatal system volume (now considered bilaterally, 0.74 mm^3^) would correspond to 0.15% of the brain volume. It is important to acknowledge that the results in this study, although corresponding to control cases which exhibit an arguably unscathed dopaminergic system (and behavior), have been designed as part of disease models [31, 34] and therefore possible differences with the wild type phenotype may exist.

Early examples for the use of the Cavalieri principle to measure the volume of individual neurons with CLSM exist [15]. Authors applied the Cavalieri principle on neurons by optically sectioning individual neurons at different depths. The current study, on the other hand, differs from the latter in that optical sectioning was not applied on individual neurons, axonal varicosities or dendrites, but on profiles of those structures as they were observed through different histological sections.

Finally, an advantage of the use of an approach based on the Cavalieri principle is that it provides robust volume estimations while relying on analysis of “two-dimensional” images. This contrasts with other stereological tools, such as the nucleator, rotator, or the optical fractionator that require fractionation on the z-axis which may be more time- and resource-consuming because it requires the analysis of tens or dozens of images per counting frame.

As mentioned, we chose to apply these methods to the description of the nigrostriatal dopaminergic system, degeneration of which is a direct cause of Parkinson Disease in humans. Because changes in axonal, dendritic and cellular structure of dopaminergic neurons accompany degeneration in humans (Fahn, 2003) and animal models, understanding its structure may help to understand the cellular mechanisms that underlie this process.

## Acknowledgements

We are grateful to Peter J. Magill, and Paul D. Dodson for generous donation of mice used in this work, to Felipe Serrano for help in illustrations, to Marcia Gaete for coaching assistance during the writing process. We are also thankful to our funding agencies, as follow: FONDECYT 1141170 and 1191497 (PH), Ring Initiative ACT1109 (FAC and PH), Geroscience Center for Brain Health and Metabolism FONDAP-15150012, FONDECYT-1190518 and MJ Fox foundation grant No. 17303 (FAC).

## Notes

### Competing Interest Statement

The authors have declared no competing interest.

## References

1. Schmitz C, Hof PR (2005) Design-based stereology in neuroscience. Neuroscience 130:813–831. 10.1016/j.neuroscience.2004.08.050

2. West MJ (2012) Estimating volume in biological structures. Cold Spring Harb Protoc 7:1129–1139. 10.1101/pdb.top071787

3. Avendaño C (2006) Stereology of neural connections: An overview. Neuroanat Tract-Tracing 3 Mol Neurons, Syst 477–529. 10.1007/0-387-28942-9_16

4. Howard C, Reed M (1998) Unbiased Stereology. Three-Dimensional Measurement in Microscopy, First. Bios Scientific Publishers - Springer, New York

5. Mouton P (2011) Unbiased Stereology: A Concise Guide, First. the Johns Hopkins University Press, Baltimore

6. Henny P, Brown MTCC, Micklem BR, et al (2014) Stereological and ultrastructural quantification of the afferent synaptome of individual neurons. Brain Struct Funct 219:631–640. 10.1007/s00429-013-0523-9

7. Faunes M, Oñate-Ponce A, Fernández-Collemann S, Henny P (2016) Excitatory and inhibitory innervation of the mouse orofacial motor nuclei: A stereological study. J Comp Neurol 524:738–758. 10.1002/cne.23862

8. Noorafshan A, Asadi-Golshan R, Abdollahifar MA, Karbalay-Doust S (2015) Protective role of curcumin against sulfiteinduced structural changes in rats’ medial prefrontal cortex. Nutr Neurosci 18:248–255. 10.1179/1476830514Y.0000000123

9. McBride JL, Pitzer MR, Boudreau RL, et al (2011) Preclinical safety of RNAi-mediated HTT suppression in the rhesus macaque as a potential therapy for Huntington’s disease. Mol Ther 19:2152–2162. 10.1038/mt.2011.219

10. Leal S, Andrade JP, Paula-Barbosa MM, Madeira MD (1998) Arcuate nucleus of the hypothalamus: Effects of age and sex. J Comp Neurol 401:65–88. 10.1002/(SICI)1096-9861(19981109)401:1<65::AID-CNE5>3.0.CO;2-D

11. Dorph-Petersen KA, Delevich KM, Marcsisin MJ, et al (2009) Pyramidal neuron number in layer 3 of primary auditory cortex of subjects with schizophrenia. Brain Res 1285:42–57. 10.1016/j.brainres.2009.06.019

12. Masilamoni GJ, Groover O, Smith Y (2017) Reduced noradrenergic innervation of ventral midbrain dopaminergic cell groups and the subthalamic nucleus in MPTP-treated parkinsonian monkeys. Neurobiol Dis 100:9–18. 10.1016/j.nbd.2016.12.025

13. García-Fiñana M, Cruz-Orive LM, Mackay CE, et al (2003) Comparison of MR imaging against physical sectioning to estimate the volume of human cerebral compartments. Neuroimage 18:505–516. 10.1016/S1053-8119(02)00021-6

14. West M, Slomianka L, Gundersen H (1991) Stereological Estimation of the Total Number of Neurons in the Subdivisions of the Rat Hippocampus Using the Optical Fractionator. Anat Rec 231:482–497

15. Prakash YS, Smithson KG, Sieck GC (1994) Application of the Cavalieri Principle in Volume Estimation Using Laser Confocal Microscopy. Neuroimage 1:325–333. 10.1006/nimg.1994.1017

16. Jacobs B, Garcia ME, Shea-Shumsky NB, et al (2018) Comparative morphology of gigantopyramidal neurons in primary motor cortex across mammals. J Comp Neurol 526:496–536. 10.1002/cne.24349

17. Tandrup T, Gundersen HJG, Jensen EBV (1997) The optical rotator. J Microsc 186:108–120. 10.1046/j.1365-2818.1997.2070765.x

18. Larsen JO (1998) Stereology of nerve cross sections. J Neurosci Methods 85:107–118. 10.1016/S0165-0270(98)00129-0

19. Bolat D, Yıldız D, Bahar S, et al (2017) A comparative study of oculomotor, trochlear and abducens nerves in Arabian foals. Biotech Histochem 92:149–156. 10.1080/10520295.2017.1288926

20. Björklund A, Dunnett SB (2007) Dopamine neuron systems in the brain: an update. Trends Neurosci 30:194–202. 10.1016/j.tins.2007.03.006

21. Matsuda W, Furuta T, Nakamura KC, et al (2009) Single nigrostriatal dopaminergic neurons form widely spread and highly dense axonal arborizations in the neostriatum. J Neurosci 29:444–453. 10.1523/JNEUROSCI.4029-08.2009

22. Gerfen CR, Surmeier DJ (2011) Modulation of striatal projection systems by dopamine. Annu Rev Neurosci 34:441–466. 10.1146/annurev-neuro-061010-113641

23. Fahn S (2003) Description of Parkinson’s Disease as a. 14:1–14

24. Fahn S (2008) The history of dopamine and levodopa in the treatment of Parkinson’s disease. Mov. Disord.

25. Deumens R, Blokland A, Prickaerts J (2002) Modeling Parkinson’s disease in rats: An evaluation of 6-OHDA lesions of the nigrostriatal pathway. Exp Neurol 175:303–317. 10.1006/exnr.2002.7891

26. Anglade P, Javoy-Agid F, AgidM Y, et al (1995) Plasticity of nerve afferents to nigrostriatal neurons in Parkinson’s disease. Ann Neurol 37:265–272. 10.1002/ana.410370219

27. Kordower JH, Olanow CW, Dodiya HB, et al (2013) Disease duration and the integrity of the nigrostriatal system in Parkinson’s disease. Brain 136:2419–2431. 10.1093/brain/awt192

28. Prasad K, Richfield EK (2008) Sporadic midbrain dopamine neuron abnormalities in laboratory mice. Neurobiol Dis. 10.1016/j.nbd.2008.07.007

29. McCormack AL, Atienza JG, Langston JW, Di Monte DA (2006) Decreased susceptibility to oxidative stress underlies the resistance of specific dopaminergic cell populations to paraquat-induced degeneration. Neuroscience. 10.1016/j.neuroscience.2006.03.069

30. Bolam JP, Pissadaki EK (2012) Living on the edge with too many mouths to feed: Why dopamine neurons die. Mov Disord. 10.1002/mds.25135

31. Janezic S, Threlfell S, Dodson PD, et al (2013) Deficits in dopaminergic transmission precede neuron loss and dysfunction in a new Parkinson model. Proc Natl Acad Sci U S A 110:. 10.1073/pnas.1309143110

32. González-Cabrera C, Meza R, Ulloa L, et al (2017) Characterization of the axon initial segment of mice substantia nigra dopaminergic neurons. J Comp Neurol 525:3529–3542. 10.1002/cne.24288

33. Meza RC, López-Jury L, Canavier CC, Henny P (2018) Role of the axon initial segment in the control of spontaneous frequency of nigral dopaminergic neurons in vivo. J Neurosci. 10.1523/JNEUROSCI.1432-17.2017

34. Oñate M, Catenaccio A, Salvadores N, et al (2020) The necroptosis machinery mediates axonal degeneration in a model of Parkinson disease. Cell Death Differ 27:1169–1185. 10.1038/s41418-019-0408-4

35. Franklin KBJ, Paxinos G (2007) The Mouse Brain in Stereotaxic Coordinates (map)

36. Conchello JA, Lichtman JW (2005) Optical sectioning microscopy. Nat Methods 2:920–931. 10.1038/nmeth815

37. Cragg SJ, Baufreton J, Xue Y, et al (2004) Synaptic release of dopamine in the subthalamic nucleus. Eur J Neurosci 20:1788–1802. 10.1111/j.1460-9568.2004.03629.x

38. Badea A, Ali-Sharief AA, Johnson GA (2007) Morphometric analysis of the C57BL/6J mouse brain. Neuroimage 37:683–693. 10.1016/j.neuroimage.2007.05.046

